# A mRNA vaccine encoding for a RBD 60-mer nanoparticle elicits neutralizing antibodies and protective immunity against the SARS-CoV-2 delta variant in transgenic K18-hACE2 mice

**DOI:** 10.1101/2022.03.21.485224

**Authors:** Pascal Brandys, Xavier Montagutelli, Irena Merenkova, Güliz T. Barut, Volker Thiel, Nicholas J. Schork, Bettina Trüeb, Laurine Conquet, Aihua Deng, Aleksandar Antanasijevic, Hyun-Ku Lee, Martine Valière, Anoop Sindhu, Gita Singh, Jens Herold

**Affiliations:** Phylex BioSciences, Del Mar, CA 92014, USA; Institut Pasteur, Université Paris Cité, Mouse Genetics Laboratory, 75015 Paris, France; Institute of Virology and Immunology, University of Bern, 3147 Mittelhäusern, Switzerland; Department of Infectious Diseases and Pathobiology, Vetsuisse Faculty, University of Bern, 3012 Bern, Switzerland; Multidisciplinary Center for Infectious Diseases, University of Bern, 3012 Bern, Switzerland; Translational Genomics Research Institute, Phoenix, AZ 85004, USA; BTS Research, San Diego, CA 92121, USA; The Scripps Research Institute, La Jolla, CA 92037, USA; Abeomics, San Diego, CA 92121, USA; Aldevron LLC, Fargo, ND 58104, USA

**Author notes:** Corresponding author: Pascal Brandys, Phylex BioSciences, Inc. 1155 Camino del Mar #502, Del Mar, CA 92014, USA.

**Keywords:** SARS-CoV-2, delta variant, mRNA vaccine, nanoparticle, receptor- binding domain, neutralizing antibody, K18-hACE2

## Abstract

Two years into the COVID-19 pandemic there is still a need for vaccines to effectively control the spread of novel SARS-CoV-2 variants and associated cases of severe disease. Here we report a messenger RNA vaccine directly encoding for a nanoparticle displaying 60 receptor binding domains (RBDs) of SARS-CoV-2 that acts as a highly effective antigen. A construct encoding the RBD of the delta variant elicits robust neutralizing antibody response with neutralizing titers an order of magnitude above currently approved mRNA vaccines. The construct also provides protective immunity against the delta variant in a widely used transgenic mouse model. We ultimately find that the proposed mRNA RBD nanoparticle-based vaccine provides a flexible platform for rapid development and will likely be of great value in combatting current and future SARS-CoV-2 variants of concern.

## INTRODUCTION

A novel zoonotic betacoronavirus that emerged in Wuhan, China at the end of 2019, subsequently named SARS-CoV-2 by the International Committee on Taxonomy of Viruses in January 2020, has resulted in the ongoing Coronavirus Disease 2019 (COVID-19) pandemic with a cumulative total of 465 million reported cases and 6 million reported deaths globally (https://covid19.who.int/table). mRNA-based vaccines for SARS-CoV-2 developed during 2020 have proven to be quite effective in preventing severe COVID-19. However, starting from the third quarter of 2020 new SARS-CoV-2 variants have repeatedly appeared and spread worldwide. As a result, the incidence rate of SARS-CoV-2 infection has increased among vaccinated individuals provided the two available mRNA vaccines (Liu et al., 2021a). From July to November 2021, the delta variant (B.1.617.2) was dominant worldwide representing over 95% of submitted sequences (https://www.epicov.org). Rapidly waning immunity of mRNA vaccines against the delta variant (Goldberg et al., 2021) prompted health authorities to recommend or mandate a third injection of legacy mRNA vaccines, with the first U.S. recommendation for a third dose of mRNA vaccine issued on August 12, 2021. The omicron variant (B.1.1.529) emerged and became dominant worldwide in less than 2 months from November 2021 to January 2022. The BA.1 sublineage initially dominant during January 2022 was rapidly replaced by the BA.1.1 sublineage during February 2022, in turn replaced by the BA.2 sublineage during March 2022, with a 75% global prevalence in submitted sequences as of March 15, 2022 (https://www.epicov.org). In the US, with a majority of the population vaccinated with at least 2 doses, daily reported new cases of infection reached a peak of 301,000 cases for the delta variant on September 7, 2021 and 1,178,000 cases for the omicron variant on January 18, 2022 (https://www.nytimes.com/interactive/2021/us/covid-cases.html). During the period of delta and omicron variant dominance, the effectiveness of 2-dose and 3- dose mRNA vaccines against emergency department/urgent care visits and hospitalizations has continuously waned over time in the US (Ferdinands et al., 2022). Two years into this global health crisis there is still a need for the global deployment of vaccines across individuals of all ages that will be effective in limiting infection and disease with current and future variants of concern (VOC).

As a result of the seriousness of the pandemic, and its complex time course, the pathobiology behind SARS-CoV-2 infection and COVID-19 illness has received considerable attention, laying the groundwork for novel diagnostic, treatment and vaccine strategies. Particularly, the SARS-CoV-2 homotrimeric spike (S) glycoprotein mediates virus entry into the host cell and comprises a N-terminal S1 surface subunit which recognizes host cell receptors, and a C-terminal S2 transmembrane subunit which promotes the fusion of the viral and cellular membranes. The receptor binding domain (RBD) of S1 binds to the host cell angiotensin-converting enzyme 2 (ACE2) receptor (Hoffmann et al., 2020; Walls et al., 2020a; Wrapp et al., 2020). The S glycoprotein is the immunogen encoded by all currently approved mRNA vaccines. The SARS-CoV-2 RBD is the target of 90% of the neutralizing activity present in COVID-19 convalescent sera and is immunodominant with multiple distinct antigenic sites (Piccoli et al., 2020). RBD-targeted neutralizing antibodies isolated from COVID-19 convalescent patients provide *in vivo* protection against SARS-CoV-2 challenge in mouse, hamster and nonhuman primates (Tortorici et al., 2020; Wu et al., 2020; Zost et al., 2020). In addition, recurrent potent neutralizing RBD-specific antibodies correlate with the plasma neutralizing activity of COVID-19 patients (Robbiani et al., 2020). All these findings indicate that the RBD is a prime target of neutralizing antibodies upon SARS-CoV-2 infection and the immunogen of choice for vaccine development.

Numerous mutations are observed throughout the genome of recent variants and these variants are highly infectious compared to the wild-type SARS-CoV-2 genome identified early in the pandemic. In most SARS-CoV-2 strain variants mutations are observed in the neutralizing antibody epitopes of the RBD, allowing escape from neutralizing antibodies (Cele et al., 2021; Collier et al., 2021; Madhi et al., 2021; Planas et al., 2021a; Wang et al., 2021). The SARS-CoV-2 delta variant is highly contagious and was rapidly spreading at the end of 2021 before the emergence of the omicron variant (Callaway, 2021). The neutralizing activity of sera from convalescent COVID-19 patients as well as sera from vaccinated individuals decreases for the delta variant compared to the wild-type (Liu et al., 2021b; Planas et al., 2021b). The RBD of the delta variant (RBDdelta) has acquired the mutations L452R and T478K that are also observed in other variants that are less infectious. It is therefore expected that the RBDdelta immunogen will elicit neutralizing antibodies directed against novel epitopes of the delta variant and other variants of interest. This effect will differ from neutralizing antibodies directed against the wild-type RBD.

The delta variant has also acquired several mutations in the N-terminal domain (NTD) of the S protein. It has been demonstrated that antibodies against a specific site on the NTD can enhance the infectivity of SARS-CoV-2 by inducing the open form of the RBD (Li et al., 2021). Furthermore, the delta variant escapes from anti-NTD neutralizing antibodies while maintaining functionally-enhancing antibody epitopes (Liu et al., 2021c), and before the emergence of the omicron variant the delta variant was likely to acquire complete resistance to wild-type spike vaccines (Liu et al., 2021d). Therefore, the RBD immunogen has an advantage in that it can avoid eliciting enhancing antibodies for the delta variant and potentially for other SARS-CoV-2 variants.

Structure-based antigen design is a widely used approach to improve immunogenicity (Graham et al., 2019; Irvine and Read, 2020; Singh, 2021) and has been applied to the RBD either by creating RBD multimers or displaying the RBD onto protein or synthetic nanoparticles (NPs) (Borriello et al. 2021, Cohen et al., 2021; Dalvie et al., 2021; He et al., 2021; King et al., 2021; Ma et al., 2020; Peachman et al., 2021; Tan et al., 2021; Walls et al., 2020b; Walsh et al., 2020). The main objective of these various designs is to increase antigen trafficking into draining lymph nodes and to promote B cell receptor clustering and B cell activation. Several RBD-NP recombinant proteins are currently evaluated in clinical trials (NCT04742738, NCT04750343, NCT05175950). In addition, many RBD-NPs designs are incompatible with mRNA vaccines, as they require various additional chemical or enzymatic reactions after protein expression, and the only RBD-based mRNA vaccine being tested in a clinical trial encodes a trimerized RBD (NCT04523571). An mRNA RBD-NP design has the advantage of combining improved immunogenicity, flexibility of design against new variants, and can be rapidly developed as a vaccine as demonstrated by first-generation mRNA vaccines.

Here, we report a messenger RNA encoding for a designed nanoparticle in which multimeric RBDdelta is displayed onto a protein scaffold composed of 60 subunits of the self-assembling bacterial protein lumazine synthase (Zhang et al., 2001).

The RBDdelta nanoparticle (RBDdelta-NP) elicits potent and protective neutralizing antibody responses in mice, with neutralizing titers an order of magnitude higher than current mRNA vaccine-elicited human sera. We further show that the mRNA RBDdelta-NP vaccine encoding for the RBDdelta protects against COVID-19 disease after SARS-CoV-2 delta variant infection in a widely used transgenic mouse model. Ultimately, our study demonstrates the utility of a mRNA vaccine directly encoding a highly immunogenic RBD nanoparticle as a flexible platform for the rapid development of effective vaccines against current and future SARS-CoV-2 variants of concern.

## RESULTS

### Design of RBD-nanoparticle

Vaccine-induced immune responses can be enhanced by NPs mimicking the size, shape, multivalency, and symmetric surface geometry of many viruses (Bachmann et al., 2021). High density display of glycosylated antigens onto protein NPs of approximately 40 nm in size enhance humoral immunity as they deposit within the B-cell follicles of the lymph nodes and generate a strong immune response (Singh, 2021). In order to display SARS-CoV-2 RBD onto a protein NP scaffold we used the self-assembling lumazine synthase (LS) from the hyper-thermophile *Aquifex aeolicus* as a protein NP scaffold (Zhang et al., 2001). The spherical LS NP scaffold consists of 60 identical subunits with strict icosahedral 532 symmetry and has been used for the display of HIV antigens (Jardine et al., 2013).

Antigens arranged in a pathogen-associated structural pattern cross-link B-cell receptors and induce the classical pathway of complement activation resulting in durable antibody responses. The optimal immune response is induced by at least 12-16 neutralizing epitopes spaced by 5-10 nm, as between repetitive immunogens on the surface of a typical RNA virus (Bachmann et al., 2021). We fused the RBD (residues 328-531) to the self-assembling LS and confirmed that with a suitable linker length, 60 copies of glycosylated RBD could be sterically accommodated. We found that such nanoparticles presenting glycosylated RBD could be secreted from mammalian (HEK 293) cells and purified by lectin chromatography with structural homogeneity, albeit with a very low yield. We further verified an optimal diameter of the NP sphere of 30 nm and an optimal spacing of 7 nm between two adjacent RBDs by negative-stain electron microscopy (Fig. 1).

**Fig 1.**
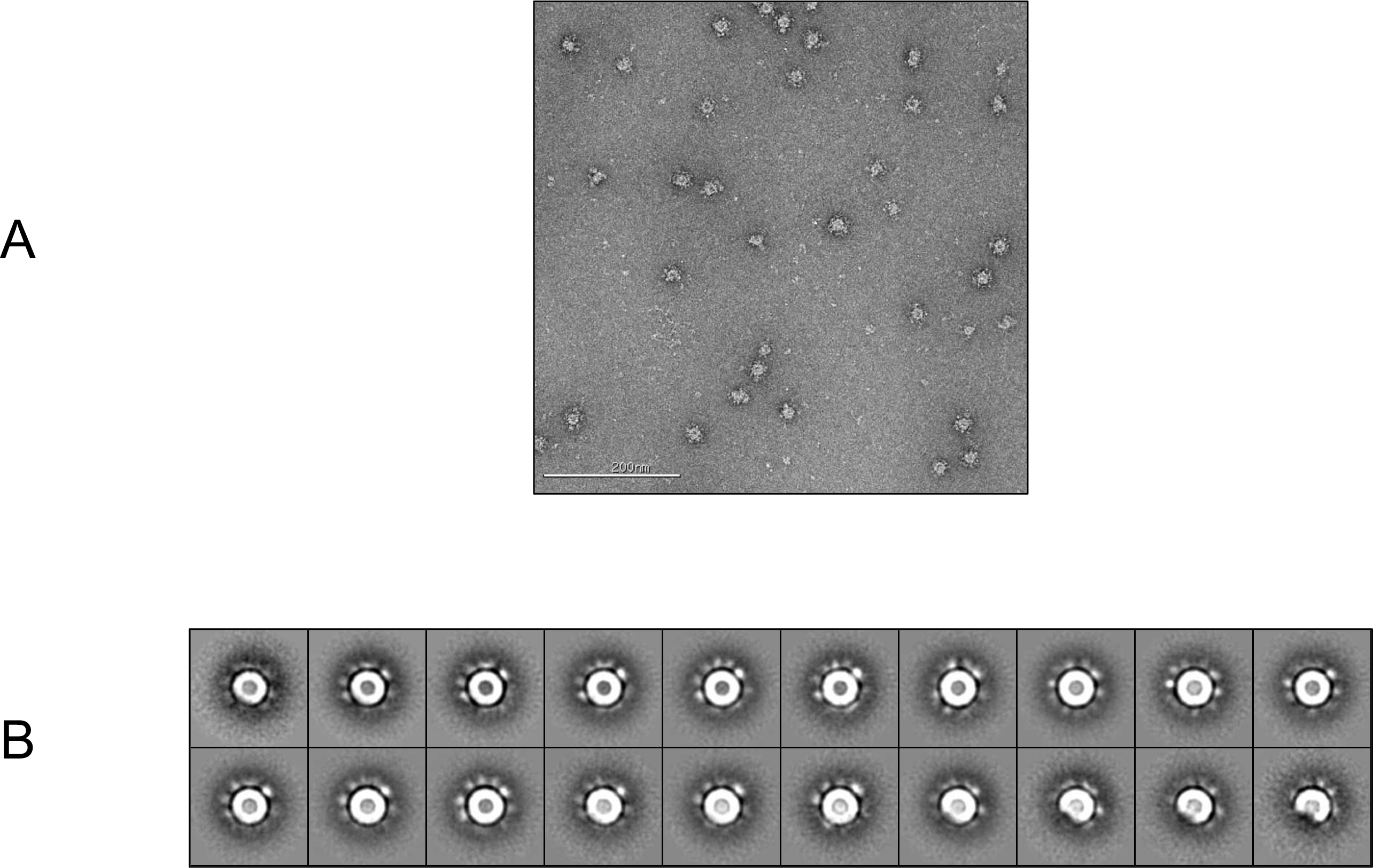
NS-EM imaging of transiently expressed RBD-NP protein A sample of glycosylated RBD-NP protein was applied to a carbon coated copper grid with uranyl-formate staining. Grids were prepared in duplicates and EM imaging was performed. **(A)** One of the 186 micrographs that were acquired. **(B)** 3271 particles were picked up manually and 2D-aligned into 20 classes. Nanoparticles are homogeneous in size, look as expected by the design with scattered RBD densities around the LS core and a diameter D ∼30 nm. Distance between adjacent RBDs on spherical NP surface is calculated as (π D^2^/60)^1/2^ ∼7 nm.

### Design of mRNA encoding for 60-mer RBD-nanoparticle

mRNA untranslated regions (UTR), including the 5’ UTR and 3’UTR, contain multiple regulatory elements and are critical for the stability and translation of mRNA into protein. For improved expression of therapeutic mRNA, a systematic evaluation of alternative UTRs is required (Asrani et al., 2018). Human beta-globin 3’ UTR was demonstrated to enhance the stability of mRNA (Adibzadeh et al., 2019), and a minimal 5’ UTR consisting of 14 nucleotides combining the T7 promoter with a Kozak consensus sequence was found to yield higher expression than human alpha globin 5’ UTR (Trepotec et al., 2019). 3’ polyadenylation with a poly(A) tail measuring 120 nucleotides compared with a shorter one and an unmasked poly(A) tail with a free 3’ end rather than one extended with unrelated nucleotides each independently enhance RNA stability and translational efficiency (Holtkamp et al., 2006). We elected to encode the poly(A) tail into the template vector rather than enzymatic polyadenylation, resulting in a limitation to about 80 nucleotides for efficient in-vitro transcription (IVT). We designed a mRNA encoding the RBD of the SARS-CoV-2 delta variant (RBDdelta) fused with LS with a minimal 5’ UTR, a human beta-globin 3’UTR and a poly(A) tail of 80 nucleotides. The mRNA was not modified, neither with guanosine/cytidine (G/C) content modification nor with chemical modification.

### Design of vector for in-vitro transcription

Overhang at the 3’ end of the poly(A) tail hampers translational efficiency of IVT RNA (Holtkamp et al., 2006), therefore we elected to introduce a site for linearization with BsmBI inserted 3’ to the poly(A) tail, resulting in a free 3’ end after the poly(A) tail. A construct with the 5’ minimal untranslated region UTR1, the human IL-2 signal sequence, a nucleotide sequence encoding RBDdelta fused with LS, the human beta-globin 3’UTR, a poly(A) tail of 80 adenosine residues and the BsmBI restriction site was cloned into a pUC19 vector.

The supercoiled pUC19 DNA was upscaled, linearized with BsmBI, and purified. In vitro transcription was performed with T7 RNA polymerase in a 2mL reaction. The mRNA was capped with a cap 1 structure on the 5’ end by vaccinia 2’-*O*- methyltransferase enzymatic capping that protects RNA from decapping and degradation (Picard-Jean et al., 2018). Capped mRNA was purified by reverse phase chromatography followed by tangential flow filtration (TFF). Final yield of purified mRNA was 2.88 g/l of IVT reaction.

### Vaccine formulation and delivery

Protamine, a small arginine-rich DNA-binding nuclear protein, condenses sperm DNA in all vertebrates (Balhorn, 2007) and is widely used to deliver different types of RNA *in vitro* and *in vivo*. Clinical grade protamine, used to condense and protect RNA from RNase degradation, shows significantly improved cytosolic delivery capacity as compared with other grades (Jarzebska, 2020). mRNA complexed with the polycationic protein protamine also results in a self-adjuvanted vaccine (Kallen et al., 2013). The mRNA was complexed by addition of clinical grade protamine to the mRNA at a mass ratio of 1:5. The vaccine was prepared on each injection day with final total mRNA concentration of 840μg/ml.

Protamine-condensed RNA was shown to be effective for vaccination using the ear pinna as injection site in mouse (Hoerr et al., 2000) and using intradermal or intramuscular needle-free injection in a human clinical trial (Alberer et al., 2017). We assessed the two injection procedures in mice and concluded at the superiority of needle-free injection for the mRNA RBD-NP vaccine (Fig. 2 A).

**Fig. 2.**
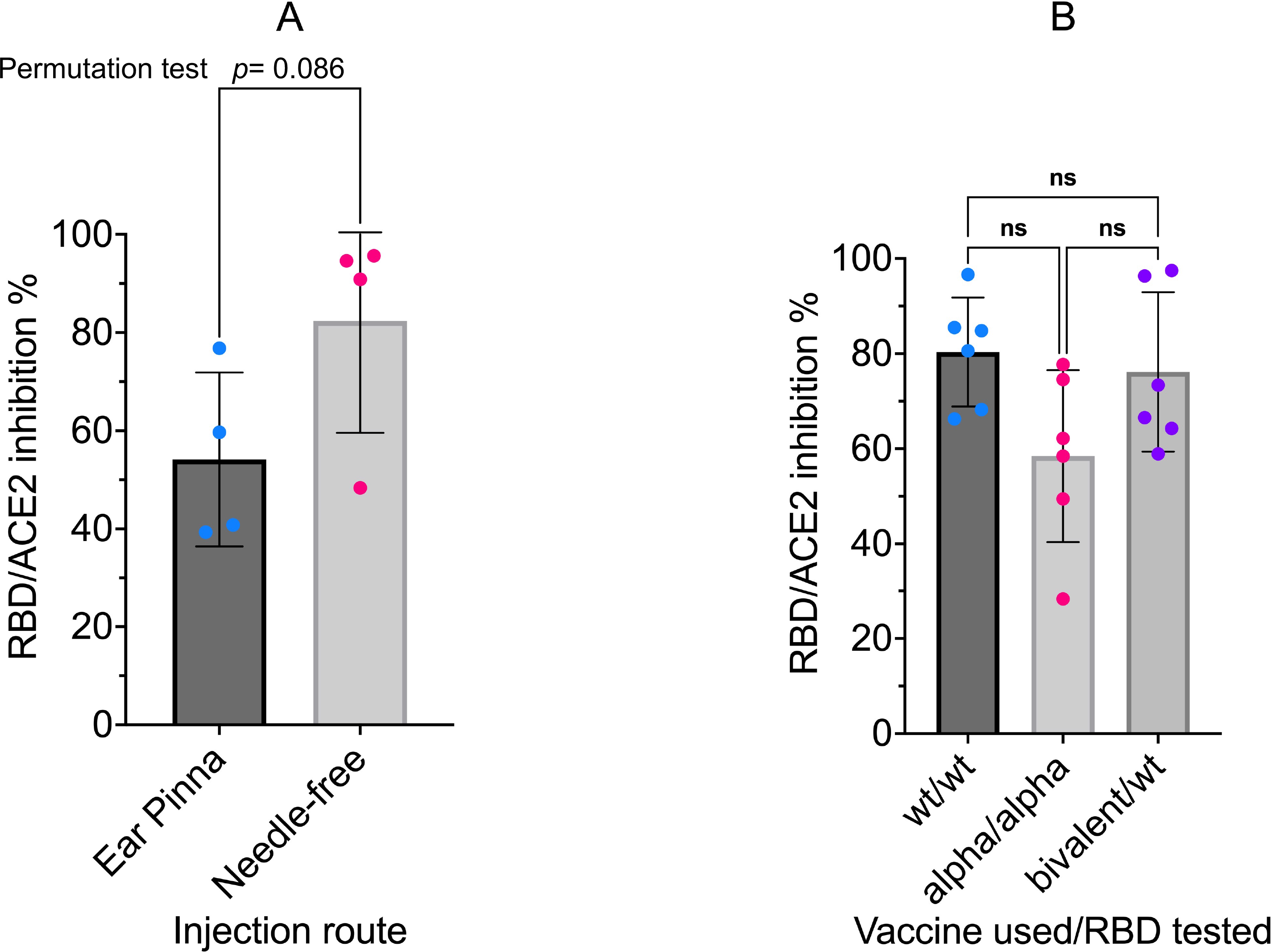
(A) Needle-free injection route is superior for RBD-NP vaccine CB6F1/J female mice 8 weeks old were primed at week 0 and boosted at week 2 with the same dose (42µg/50µl) of the mRNA RBD-NP vaccine with the wild type RBD. Group 1 (n= 4) by i.d. injection at the ear pinna and Group 2 (n= 4) by i.m. needle-free injection at the caudal thigh with the modified Tropis® injector. Serum samples collected in week 4 were analyzed with the SARS-CoV-2 neutralization antibody assay. Group 1: mean= 54.2%, SEM= 8.8, Group 2: mean= 82.4%, SEM= 11.4. The difference in means between the two samples of small size was analyzed with a standard two-sided permutation test (*p*-value= 0.0857). (B) **Anti-RBD neutralizing antibodies compared in sera induced by monovalent and bivalent mRNA RBD-NP vaccines** CB6F1/J female mice 8 weeks old were primed at week 0 and boosted at week 3 by needle-free i.m. injection at the caudal thigh with the modified Tropis® injector and the same dose (42µg/50µl) of three different mRNA RBD-NP vaccines. Group 1 (n= 6) received the monovalent mRNA RBDwt-NP vaccine with wild type RBD (RBDwt). Group 2 (n= 6) received the monovalent mRNA RBDalpha-NP vaccine with alpha variant RBD (RBDalpha). Group 3 (n= 6) received the bivalent mRNA vaccine with equal dose of mRNA RBDwt-NP and mRNA RBDalpha-NP. Serum samples collected in week 6 were analyzed with the SARS-CoV-2 neutralization antibody assay with different HRP conjugated RBDs. G1 was tested for RBDwt-ACE2 interaction, G2 was tested for RBDalpha-ACE2 interaction and G3 was tested for RBDwt-ACE2 interaction. *p* values in one-way ANOVA multiple comparison: wt/wt vs. bivalent/wt = 0.89; wt/wt vs. alpha/alpha = 0.07; alpha/alpha vs. bivalent/wt = 0.16. No significant difference among the means of monovalent and bivalent vaccines for inhibition of RBD-ACE2 interaction by anti-RBD neutralizing antibodies.

### Immunization with mRNA RBDdelta-NP vaccine elicits high serum anti- RBDdelta antibody titers and SARS-CoV-2 delta neutralizing titers in mice

To investigate the prophylactic potential of the mRNA RBDdelta-NP vaccine against the delta variant of SARS-CoV-2 we gave to one group (n=8) of mice a prime/boost regimen of the mRNA RBDdelta-NP vaccine. CB6F1/J female mice 8 weeks old were primed at week 0 and boosted at week 3 with the same dose of 42µg/50µl by intramuscular injection at the caudal thigh with a needle-free injection system (Tropis® injector modified for mouse injection, PharmaJet). Blood was collected and serum prepared on weeks 0 (prior to prime), 3 (prior to boost) and 6.

To determine if the mRNA RBDdelta-NP vaccine elicited SARS-CoV-2 delta neutralizing antibodies we analyzed samples of week 0, 3 and 6 with the cPass™ SARS-CoV-2 neutralization antibody detection kit (GenScript, Cat #L00847). The assay detected any antibodies in serum that neutralize the interaction between the RBDdelta and the ACE2 receptor. The RBDdelta-ACE2 interaction inhibition rate was calculated with the net optical density (OD) of sample and negative control as Inhibition = (1 – OD value of sample/OD value of negative control) x 100%. Neutralizing antibodies were detected with a cutoff inhibition value of 30% established in a human clinical study.

In all 8 mice receiving the mRNA RBDdelta-NP vaccine for all 8 week 3 samples and all 8 week 6 samples SARS-CoV-2 delta neutralizing antibodies were detected in the mouse sera. For all random week 0 samples no neutralizing antibodies were detected. High inhibition at week 3 (mean 87.0%, standard deviation 10.3%) and very high inhibition at week 6 (mean 98.0%, standard deviation 0.6%) was observed in all mouse sera. There was a significant difference between week 3 and 6 (Student t-test, *p-*value= 0.01) (Fig. 3 A).

**Fig. 3.**
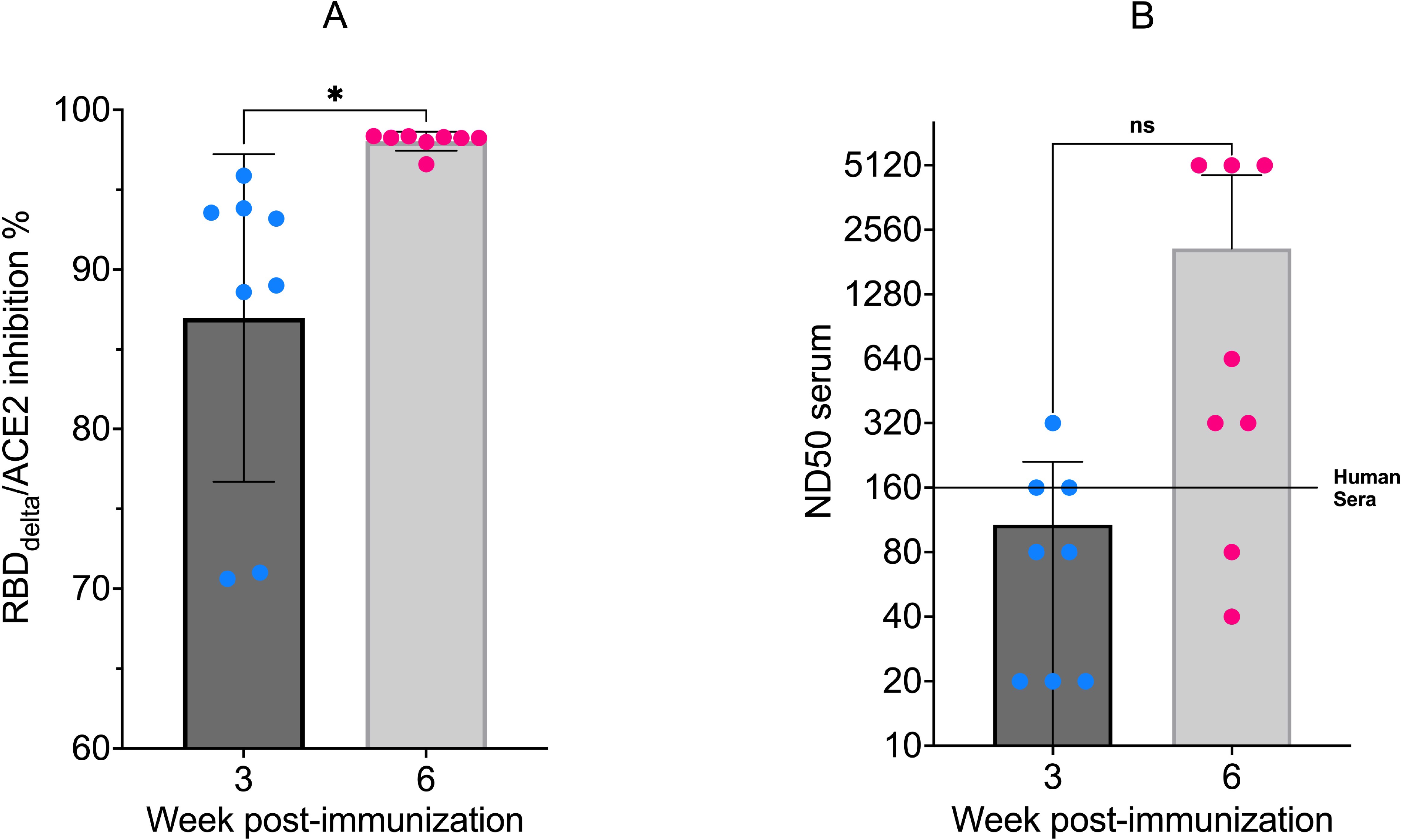
(A) RBDdelta-ACE2 interaction inhibition by neutralizing antibodies in mice. Sera samples analyzed with the SARS-CoV-2 neutralization antibody detection assay. RBDdelta-ACE2 interaction inhibition rate at week 3 and 6 in CB6F1/J female mice (n= 8). Week 3 and 6 were significantly different in two-tailed paired t-test (*p*-value= 0.016). (B) **SARS-CoV-2 delta variant neutralization in mice** Sera samples analyzed with the SARS-CoV-2 delta variant neutralization test. Median neutralization dose (ND50) at week 3 and 6 in CB6F1/J female mice (n= 8). BNT162b2-elicited human sera as reference. Week 3 and 6 were not significantly different in two-tailed paired t-test (*p*-value= 0.063).

The sera of all mRNA RBDdelta-NP-immunized mice collected at week 3 and 6 were also tested for neutralization of cell infection by authentic SARS-CoV-2 delta coronavirus. Two sera of human patients vaccinated with a regular vaccination schedule of BNT162b2 (Pfizer-BioNTech) mRNA vaccine and collected three weeks after the second dose were used as positive controls. The mRNA RBDdelta- NP needle-free intramuscular vaccination schedule induced effective neutralizing antibodies against the SARS-CoV-2 delta variant in 5 mouse sera at week 3 and in all 8 mouse sera at week 6, with median neutralization dose (ND50) ranging from 40 to 5120 at week 6. Mean ND50 was 108 at week 3 and 2095 at week 6, whereas mean ND50 of positive controls was 160. Thus, mean ND50 of sera 3 weeks after the second dose was 13-fold as compared with BNT162b2-elicited human sera. For 3 mouse sera ND50 at week 6 was 32-fold as compared with BNT162b2-elicited human sera (Fig. 3 B).

### Immunization with mRNA RBDdelta-NP vaccine protects transgenic K18- hACE2 mice from SARS-CoV-2 delta variant challenge and elicits neutralizing antibodies

Transgenic mice expressing human angiotensin-converting enzyme 2 under the control of human cytokeratin 18 promoter (K18-hACE2) are highly susceptible to SARS-CoV-2 infection and represent a suitable animal model for the study of viral pathogenesis, and for identification and characterization of prophylactic vaccines for SARS-CoV-2 infection and associated severe COVID-19 disease (Oladunni et al., 2020).

To assess the prophylactic effect of the mRNA RBDdelta-NP vaccine against the delta variant of SARS-CoV-2 we gave to one group (n=16) of K18-hACE2 transgenic mice a prime/boost regimen of the mRNA RBDdelta-NP vaccine (Fig. 4). Female K18-hACE2 transgenic mice at 8 weeks old were primed at week 0 and boosted at week 3 with the same dose of 42µg/50µl by intramuscular injection at the caudal thigh with the needle-free injection system. Blood was collected and serum prepared in week 0 (prior to prime), 3 (prior to boost), 6 and 8 (prior to viral challenge). A second control group (n=16) of female K18-hACE2 transgenic mice of same age did not receive any vaccine. Both groups were housed under the same conditions, including a 3-day international shipment during week 4 from California (after immunization) to France (before viral challenge).

**Fig. 4.**
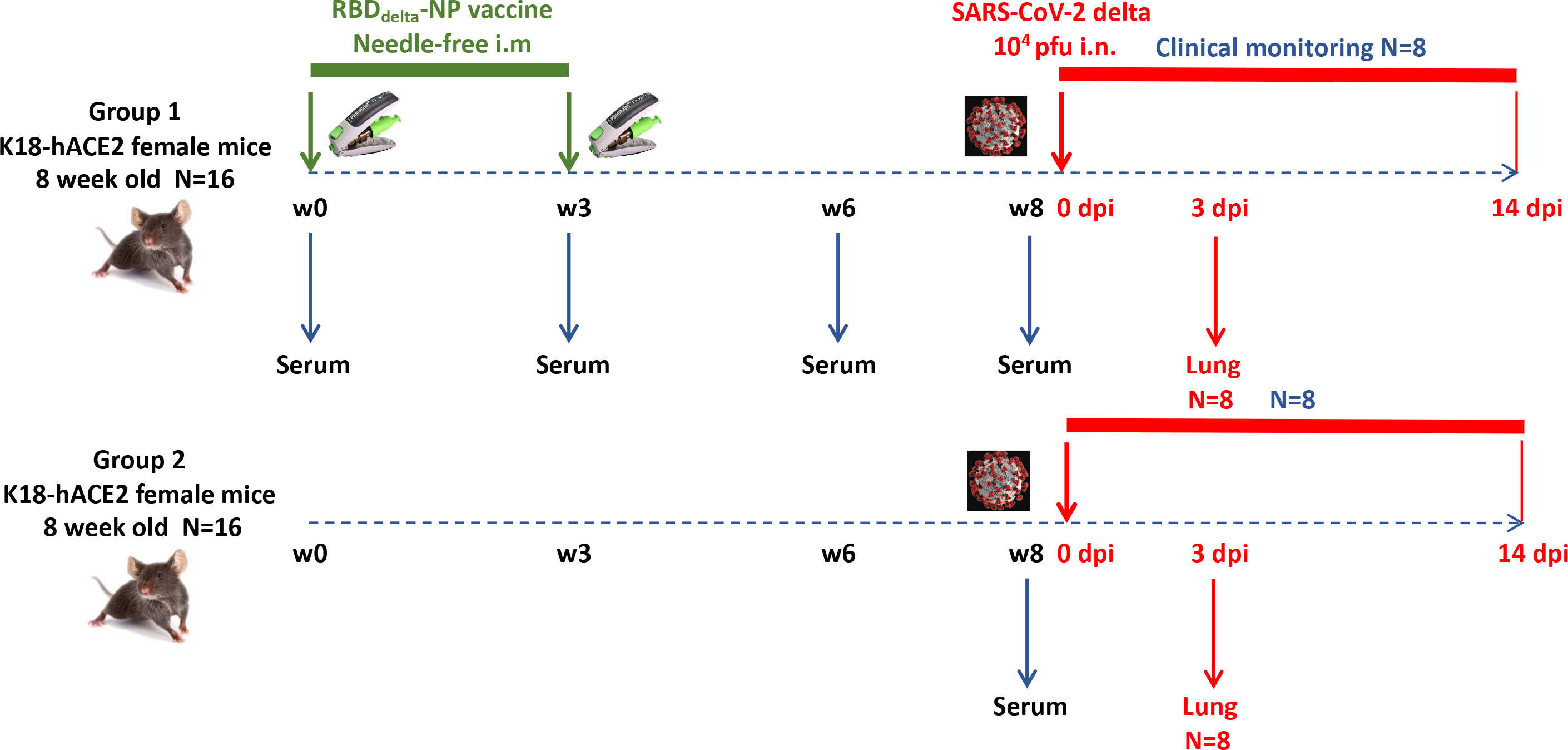
Experimental design of immunogenicity and protection efficacy study of mRNA RBDdelta-NP vaccine in K18-hACE2 transgenic mice Two groups (n=16): Group 1 immunized with mRNA RBDdelta-NP vaccine, Group 2 unvaccinated control. In each group 8 mice are sacrificed at 3 dpi for lung collection and 8 mice are followed for clinical signs until 14 dpi.

To determine if mRNA RBDdelta-NP vaccine elicited neutralizing antibodies in mouse sera, we analyzed samples of week 0, 3, 6 and 8 with the cPass™ SARS- CoV-2 neutralization antibody detection assay. In the 16 mice receiving the mRNA RBDdelta-NP vaccine regimen, SARS-CoV-2 delta neutralizing antibodies were detected in the mouse sera of 14/16 week 3 samples, 16/16 week 6 samples and 16/16 week 8 samples. No neutralizing antibodies were detected in random week 0 samples. There was no significant difference between week 6 and week 8 for RBDdelta-ACE2 interaction inhibition rates. In 12 mice there was strong inhibition rate at week 6 (mean 89.6%, standard deviation 5.4%) and week 8 (mean 90.6%, standard deviation 4.0%) (Fig. 5 A and B).

**Fig. 5.**
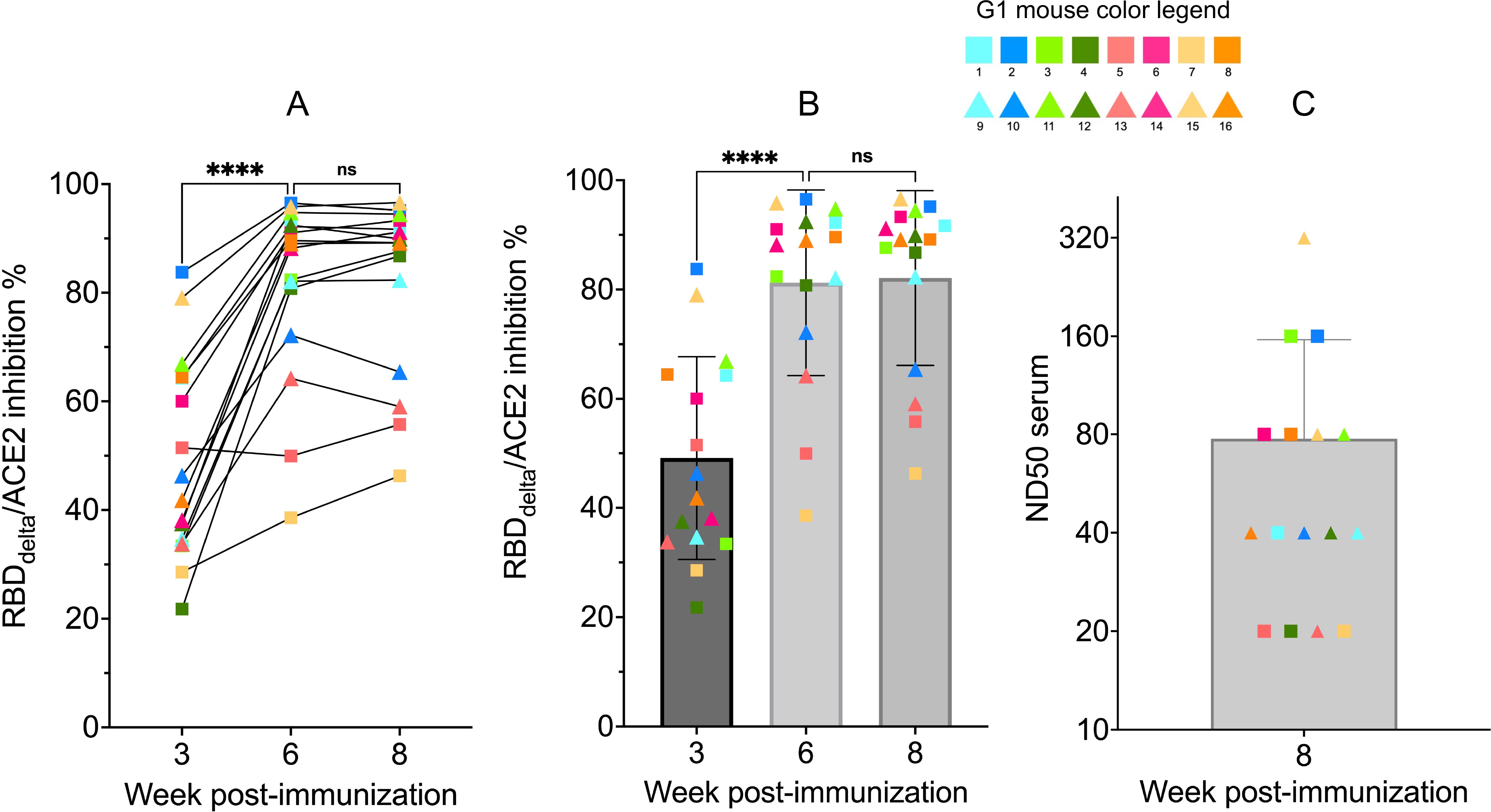
RBDdelta-ACE2 interaction inhibition by neutralizing antibodies and SARS-CoV-2 delta variant neutralization in K18-hACE2 transgenic mice Sera samples of immunized K18-hACE2 transgenic female mice (n= 16) analyzed with the SARS-CoV-2 neutralization antibody assay and the SARS-CoV-2 delta variant neutralization test (Group 1 with individual color code, squares indicate mice euthanized at 3 dpi and triangles indicate mice followed until 14 dpi). **(A)** RBDdelta-ACE2 interaction inhibition rate at week 3, 6 and 8 with paired individual values. **(B)** RBDdelta-ACE2 interaction inhibition rate at week 3, 6 and 8. Week 3 and 6 were significantly different in one-way ANOVA multiple comparison (*p*- value< 0.0001). Week 6 and 8 were not significantly different (*p*-value= 0.66). **(C)** Median neutralization dose (ND50) at week 8.

The sera of all 16 RBDdelta-NP-immunized mice collected at week 8 prior to viral challenge was also tested for neutralization of cell infection by authentic SARS-CoV-2 delta coronavirus as above. ND50 ranged from 20 to 320 with a mean of 78, whereas ND50 of human serum positive control was 80 (Fig. 5 C). As expected ND50 values after Log2 transformation correlated significantly with the RBDdelta- ACE2 interaction inhibition rates of the 16 week 8 sera samples (non-parametric Spearman test: correlation coefficient r= 0.79, *p-*value= 0.0005). A comparison of normalized results between the two experiments showed a mean ND50 in CB6F1/J mice 1.3 log10 above the mean in K18-hACE2 transgenic mice.

At week 8 all 32 transgenic mice were inoculated with an intranasal dose of 10^4^ plaque-forming units (PFU) of SARS-CoV-2 delta variant (B.1.617.2 strain hCoV- 19/France/HDF-IPP11602i/2021). Eight immunized mice and 8 control mice were followed for 14 days post-infection (dpi) for body weight variation and clinical signs of COVID-19 disease (Fig. 4). All 8 control mice showed signs of severe disease with rapid body weight loss and reached termination criteria at 7 or 8 dpi (Fig. 6 and 7). All 8 immunized mice showed body weight gain at 14 dpi with 3 mice experiencing a mild symptomatic episode at 5-6 dpi followed by rapid recovery at 7-8 dpi (Fig. 6 and 7).

**Fig. 6.**
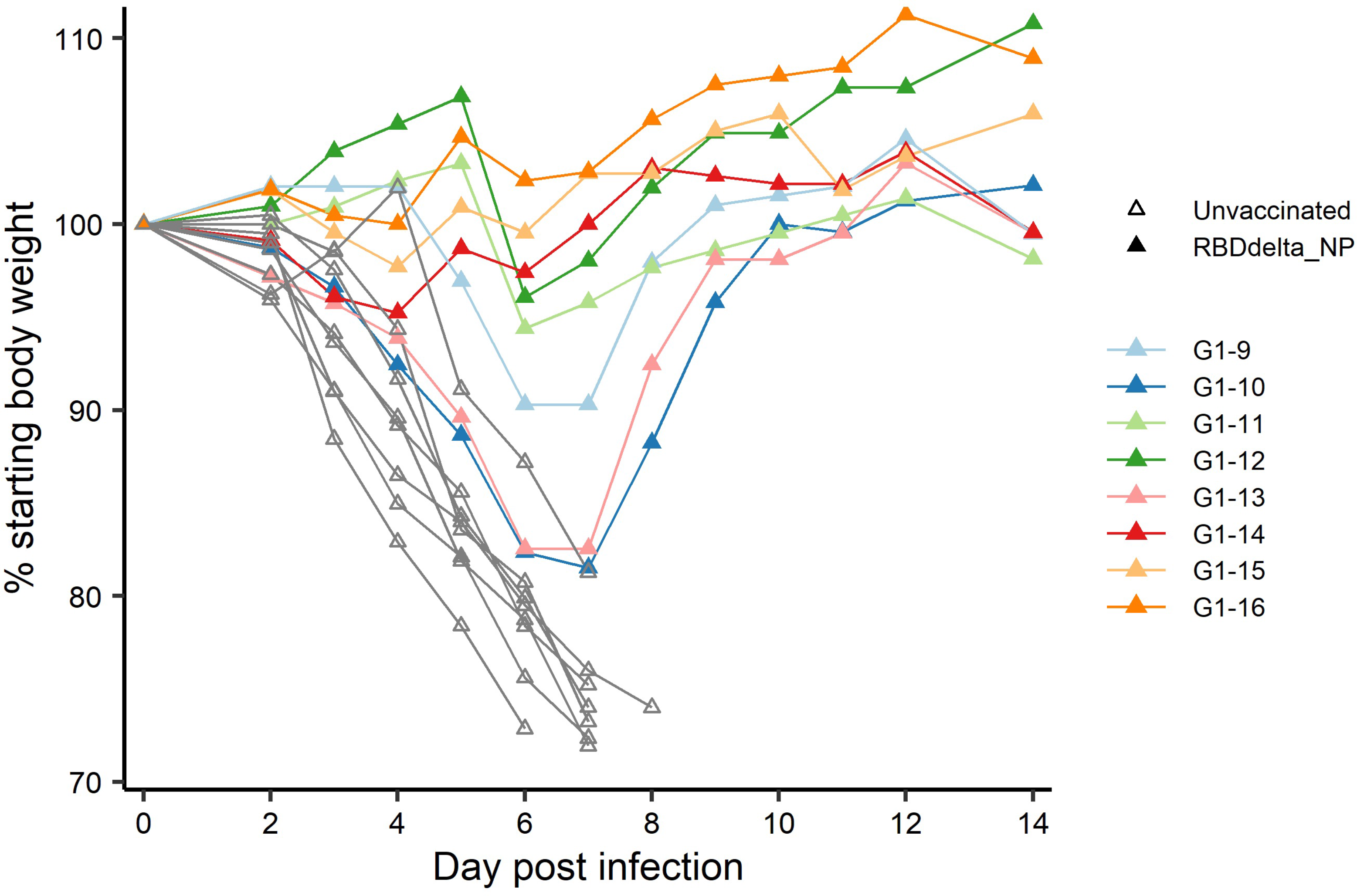
Individual body weight 0-14 dpi Daily individual weights of infected K18-hACE2 transgenic mice expressed as percentage of starting body weight. Immunized Group 1 (n= 8) is with individual color code and unvaccinated Group 2 (n= 8) is in grey.

**Fig. 7.**
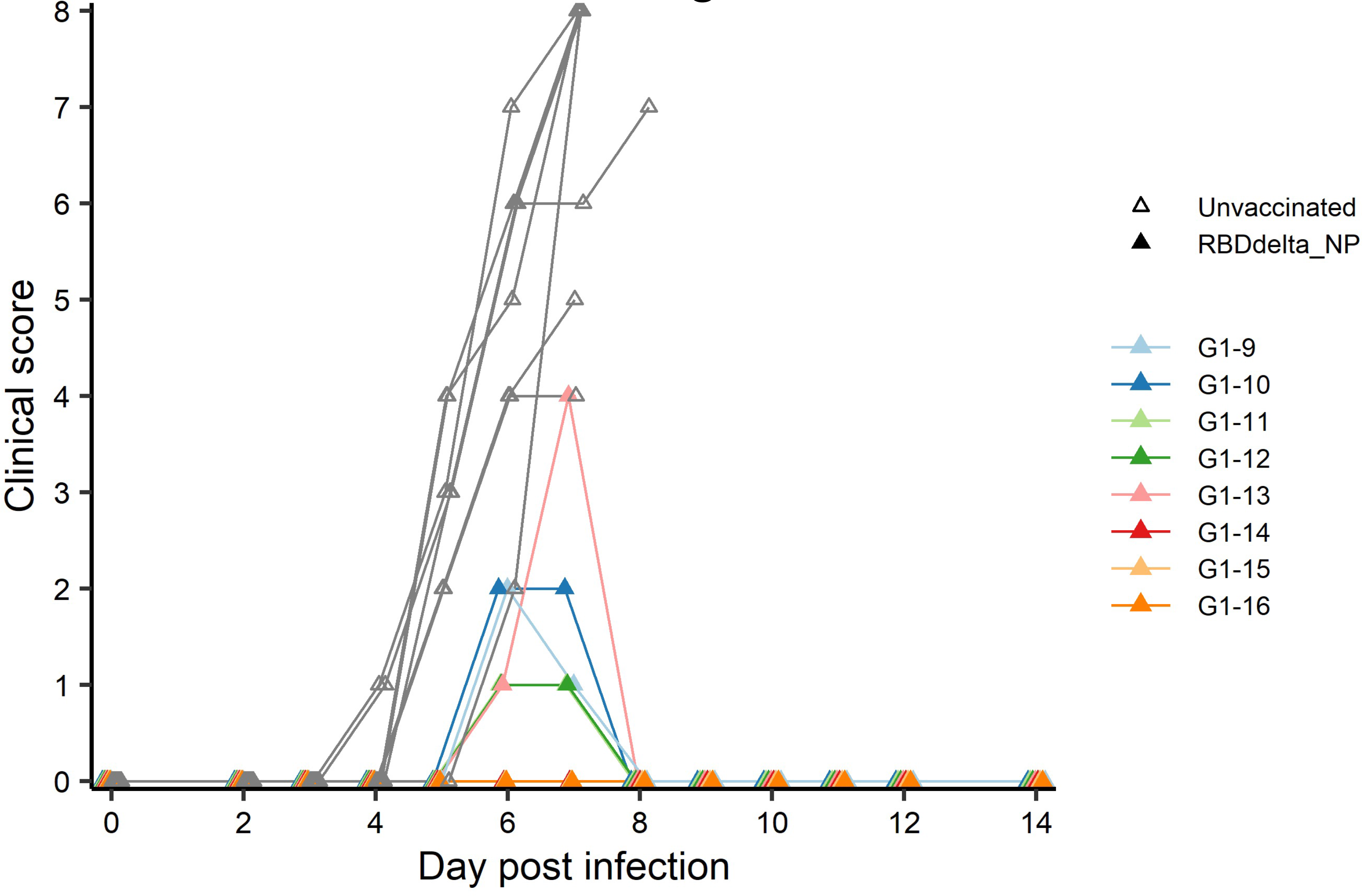
Individual clinical score 0-14 dpi Daily individual clinical score of infected K18-hACE2 transgenic mice. Each of four criteria (ruffled fur, hunched posture, reduced locomotion and difficult breathing) received a 0-3 score and were added into a global score. Immunized Group 1 (n= 8) is with individual color code and unvaccinated Group 2 (n= 8) is in grey.

Eight immunized mice and 8 unvaccinated controls were euthanized at 3 dpi for measurement of viral load and viral titer in lung homogenates (Fig. 4). Viral load showed a significant reduction in immunized mice (Student t-test, *p*= 0.0088) (Fig. 8 A). Viral titration on Vero-E6 cells showed a more significant reduction in immunized mice (Student t-test, *p*= 0.0015) and half of the immunized mice showed high protection with mean viral titers 2.6 log10 below the mean of control mice (Fig. 8 B).

**Fig. 8.**
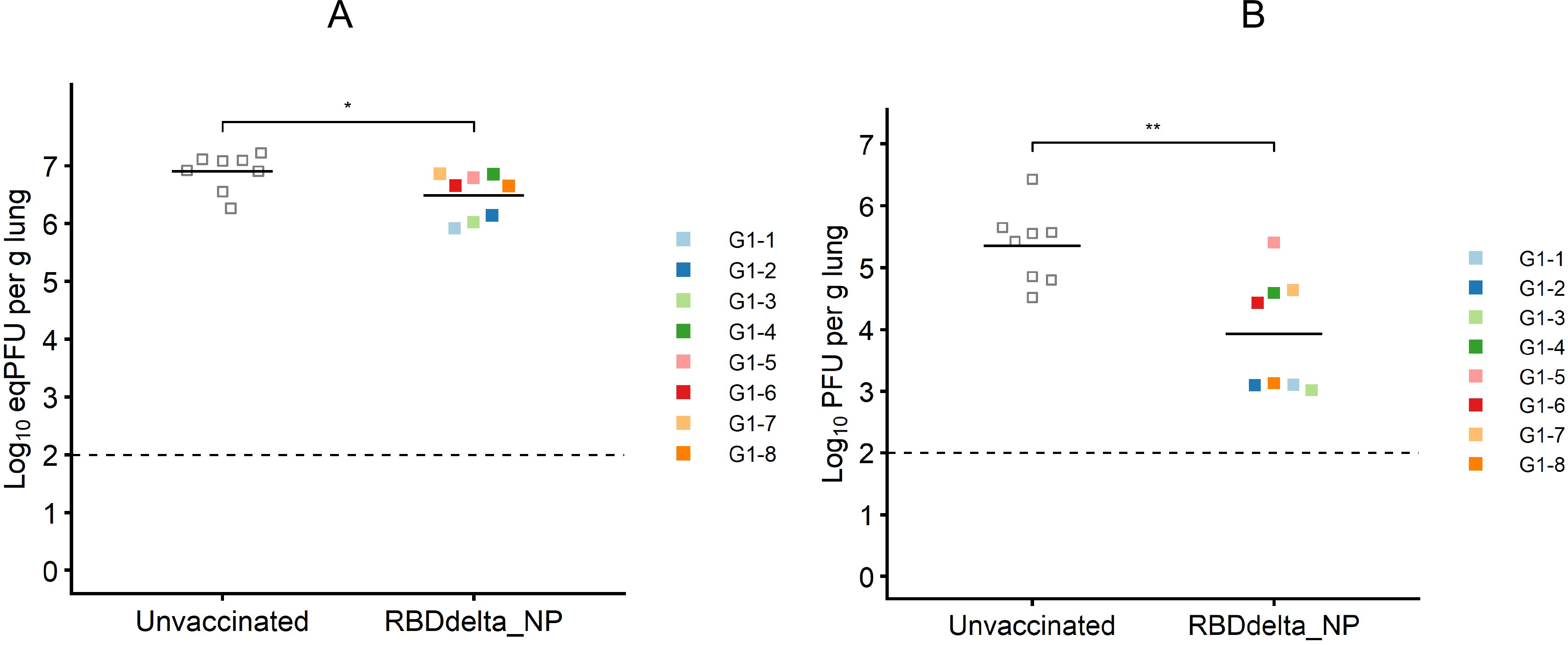
Protection against lung infection at 3 dpi On day 3 post-infection K18-hACE2 transgenic mice were sacrificed and lungs were collected to assess viral titers. Immunized Group 1 (n= 8) is with individual color code and unvaccinated Group 2 (n= 8) is in grey. **(A)** Viral load in lungs expressed in eqPFU per gram of lung tissue as determined by RT-qPCR. **(B)** Viral titer in lungs in plaque assay expressed in log10PFU per gram of lung tissue.

### Elevated *Ccl5* and *Il12* expression indicates an adaptive cell-mediated immune response to mRNA RBDdelta-NP vaccine

SARS-CoV-2 infection of K18-hACE2 transgenic mice results in the activation of cytokines and chemokines, as described during the development of severe COVID-19 disease in humans (Oladunni et al., 2020). We also assessed the variation of ten inflammatory cytokine and chemokines in the RNA extracted from total lung homogenates at 3 dpi (Fig. 9 A). We found that reduced expression of C-C motif ligand 2 (*Ccl2*), C-C motif ligand 3 (*Ccl3*) and C-X-C motif ligand 10 (*Cxcl10*) contents in immunized mice significantly correlated with reduced lung infection expressed in log10PFU per gram of lung (Fig. 9 B). Therefore, the mRNA RBDdelta-NP vaccine also confers protection against lung inflammation mediated by SARS-CoV-2 delta variant infection, as evaluated by titration. We also found a significant increase of the relative expression of subunit p40 of interleukin 12 (*Il12p40*) and chemokine C-C motif ligand 5 (*Ccl5*) in lung homogenates of 8 immunized mice versus 8 unvaccinated and challenged mice (Fig. 9 A). *Ccl5* expression is induced during the later stage of inflammation and is an intrinsic property acquired by CD8^+^ memory T cells (Marçais et al., 2006) and *Ccl5* mRNA is translationally silenced in memory phenotype CD8^+^ T cells until T cell receptors (TCR) stimulation (Swanson et al., 2002). In mouse viral lung disease *Ccl5* mRNA in the lung peaks less than a day after primary viral infection with *Ccl5* production by resident epithelial cells and lung macrophages during the innate immune response, and then in a second wave at 7 dpi, primarily with *Ccl5* production by infiltrating CD4^+^ and CD8^+^ T cells (Culley at al., 2006). Therefore, elevated *Ccl5* expression in immunized mice at 3 dpi was indicative of sustained numbers of memory phenotype T cells in the lung tissue. Furthermore, interleukin 12 (*Il12*) expression during infection determines the type and duration of adaptive immune response. *Il12* promotes the differentiation of naïve CD4^+^ T cells into T helper 1 (Th1) cells and aids T-cell expansion and proliferation in cell-mediated immunity (Watford, 2003). Therefore, the combined increase of *Ccl5* and *Il12* expression at 3 dpi was indicative of long-term adaptive cell-mediated immune response induced by the mRNA RBDdelta-NP vaccine and warrants further investigation of virus-specific T and B cell response.

**Fig. 9.**
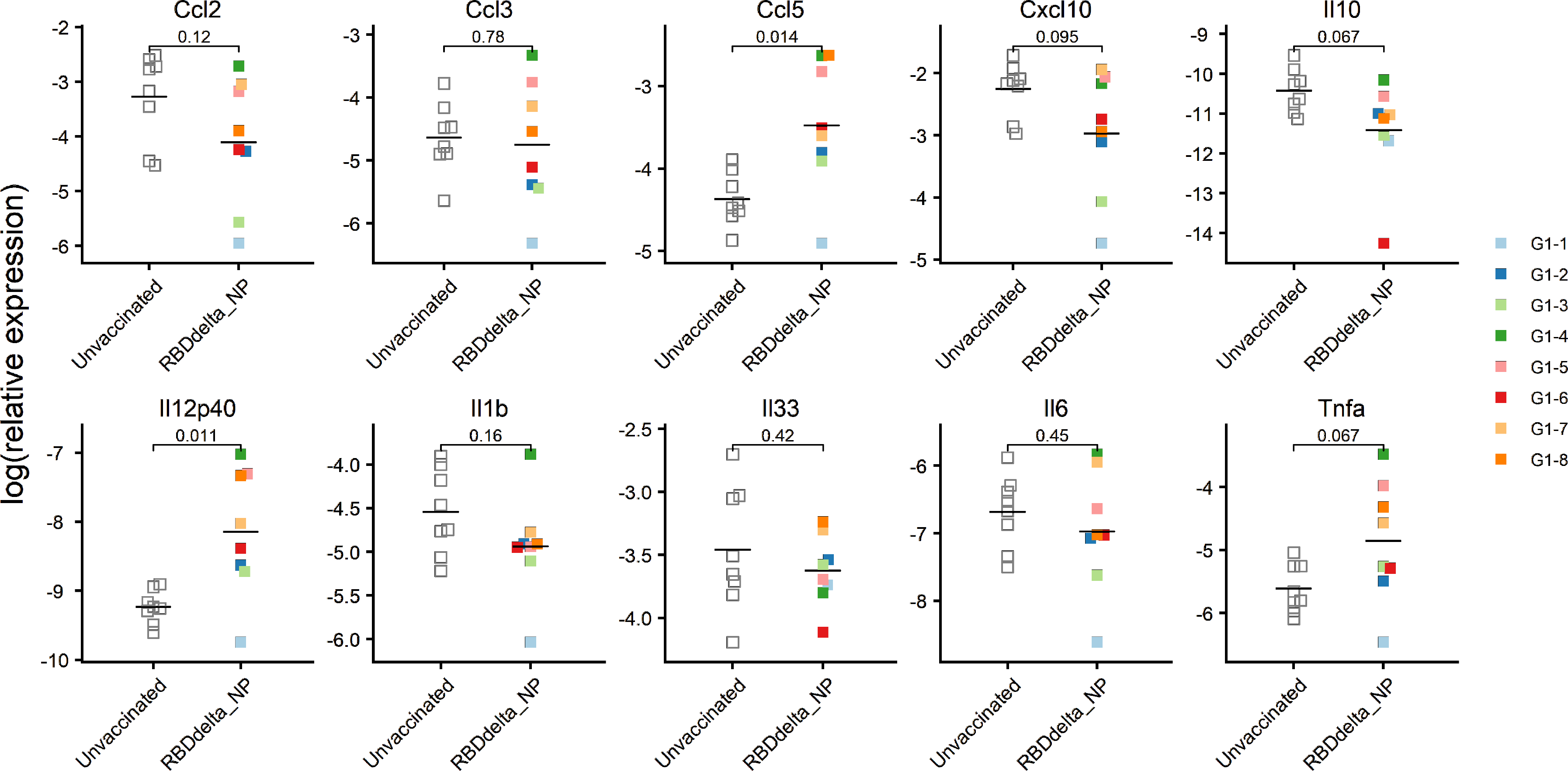
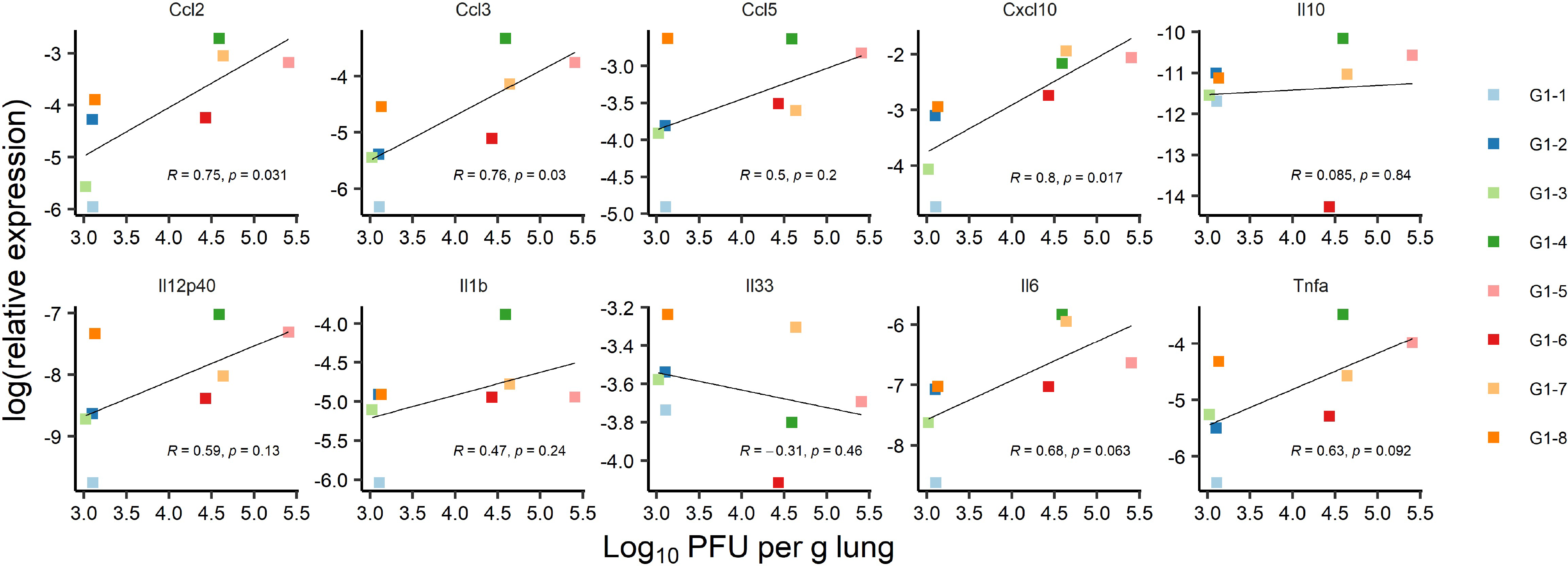
mRNA expression of ten cytokines and chemokines in the lungs at 3 dpi **(A)** Relative expression in log2 of each cytokine or chemokine mRNA in lung homogenates of infected K18-hACE2 transgenic mice as determined by RT- qPCR. Immunized Group 1 (n= 8) is with individual color code and unvaccinated Group 2 (n= 8) is in white. Expression of Il12p40 (*p*-value= 0.011) and Ccl5 (*p*- value= 0.014) is significantly increased. **(B)** Correlation between lung infection and cytokine or chemokine expression was assessed in infected K18-hACE2 transgenic mice of immunized Group 1 sacrificed at 3 dpi for lung collection (n= 8 with individual color code). Simple linear regression between log10PFU per gram of lung tissue at 3 dpi and log2 of relative expression for each cytokine or chemokine. Correlation is significant for Ccl2 (*p*-value= 0.031), Ccl3 (*p*-value= 0.03), and Cxcl10 (*p*-value= 0.017) chemokines, with correlation coefficients ranging from 0.75 to 0.8.

### Protection against SARS-CoV-2 delta infection and associated COVID-19 disease correlates with anti-RBDdelta antibody titers in K18-ACE2 transgenic mice

To determine the prophylactic effect of immune response against the RBDdelta-NP antigen we assessed the correlation between clinical results and serology results in vaccinated mice. RBDdelta-ACE2 inhibition ratio in mouse sera showed significant correlation, as measured by linear regression analysis and ranked by decreasing *p-*value, with body weight variation (Fig. 10 A) and clinical score (Fig. 10 B). Thus, anti-RBDdelta antibodies elicited by the mRNA RBDdelta-NP vaccine are an important correlate of protection against COVID-19 disease in K18-ACE2 transgenic mice.

**Fig. 10.**
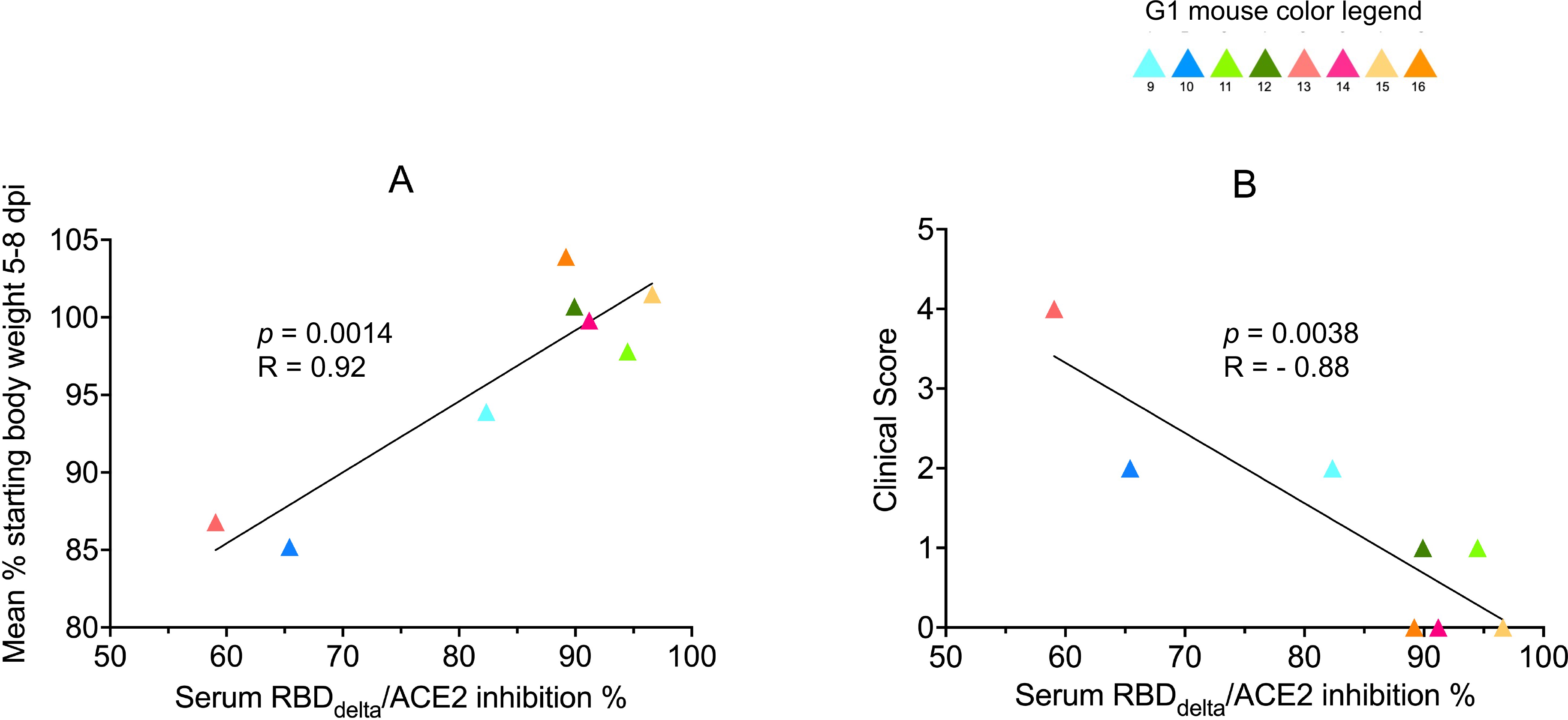
Protection against COVID-19 disease correlates with RBDdelta-ACE2 interaction inhibition by neutralizing antibodies Correlation between SARS-CoV-2 neutralization antibody assay results and clinical results was assessed in infected K18-hACE2 transgenic mice of immunized Group 1 followed until 14 dpi (n= 8 with individual color code). **(A)** Simple linear regression between RBDdelta-ACE2 interaction inhibition rate at week 8 and body weight variation between day 0 and mean of body weights during acute disease period (5-8 dpi). Correlation is significant (*p*-value= 0.0014, R= 0.92). **(B)** Simple linear regression between RBDdelta-ACE2 interaction inhibition rate at week 8 and highest reached clinical score. Correlation is significant (*p*-value= 0.0038, R= -0.88).

We also assessed the correlation between infection in the lungs and serology results. The correlation with RBDdelta-ACE2 inhibition ratio (Fig. 11 A) was not significant. However, viral titer in lungs after infection correlated significantly with serum neutralization of cell infection with SARS-CoV-2 delta variant (*p-*value=

**Fig. 11.**
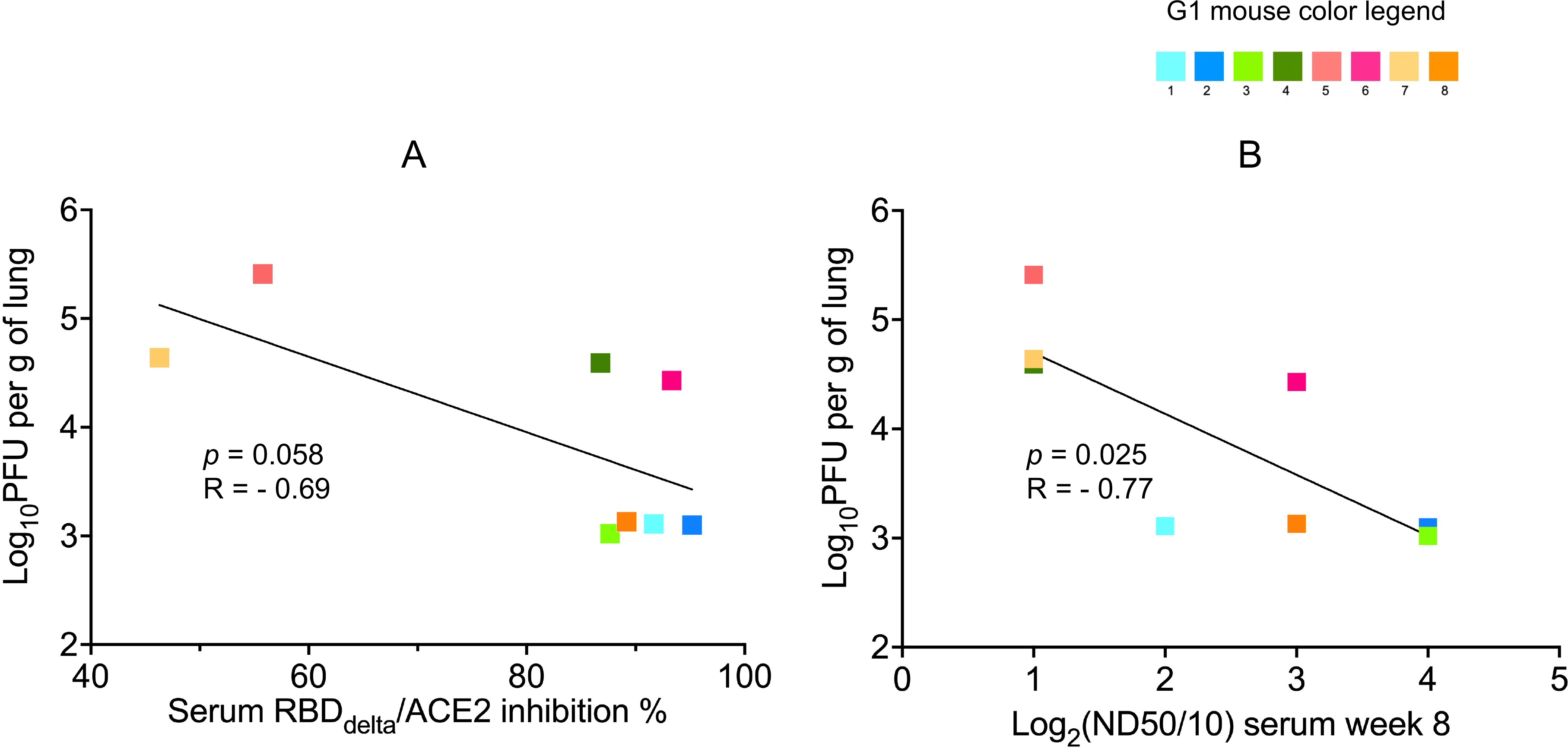
Protection against lung infection with SARS-Cov-2 delta variant correlates with serum SARS-CoV-2 delta variant neutralization Correlation between results of SARS-CoV-2 delta variant neutralization test, results of neutralization antibody assay, and lung infection was assessed in infected K18-hACE2 transgenic mice of immunized Group 1 sacrificed at 3 dpi for lung collection (n= 8 with individual color code). **(A)** Simple linear regression between RBDdelta-ACE2 interaction inhibition rate at week 8 and log10PFU per gram of lung tissue at 3 dpi. Correlation is not significant (*p*-value= 0.058, R= - 0.69) and non-parametric Spearman correlation is also not significant (p-value= 0.083, R= -0.67). **(B)** Simple linear regression between log2(ND50/10) (or number of dilutions after initial 1:10 dilution) analyzed with the SARS-CoV-2 delta variant neutralization test at week 8 and log10PFU per gram of lung tissue at 3 dpi. Correlation is significant (*p*-value= 0.025, R= -0.77) and non-parametric Spearman correlation is more significant (p-value= 0.006, R= -0.89).

0.025 and *p*-value= 0.006 in non-parametric Spearman correlation) (Fig. 11 B).

## DISCUSSION

In this study we demonstrated that mRNA RBDdelta-NP vaccine elicits a robust neutralizing anti-RBDdelta antibody response with high serum neutralizing titers that are an order of magnitude above sera elicited by current mRNA vaccination regimen. We also showed that a prime/boost of mRNA RBDdelta-NP vaccine effectively protects against the SARS-CoV-2 delta variant infection and associated COVID-19 disease in a widely used transgenic mouse model. We further showed that protection against lung infection and COVID-19 disease is correlated with serum anti-RBDdelta antibody titers. High protection against lung infection in 4/8 mice, and complete protection against COVID-19 disease in 3/8 mice, was observed with high mean values for RBDdelta-ACE2 interaction inhibition rate (91.5%) and median neutralization dose (126) in the mouse sera samples. We finally showed increased expression of CCL5 and IL-12 in the lungs indicative of long-term cell-mediated immune response induced by the mRNA RBDdelta-NP vaccine. This result calls for demonstration of virus-specific T and B cell response in future studies with RBD-NP vaccines.

We primarily attribute the variability of vaccine responsiveness between immunized mice to the needle-free injection procedure. The Tropis® injector was designed for humans with a large syringe head around the syringe nozzle which is inadequate for small animals. The injector was adapted for mice with a prototype delivering half the volume under reduced pressure, but the procedure with the prototype results in loss of material during injection and a mix of intradermal and intramuscular injection in mice. Similar experiments in larger animal models will likely result in reduced variability.

The mRNA RBD-NP vaccine platform can readily be used for the design of multivalent mRNA vaccines encoding for different RBD variants displayed onto the LS NP scaffold. Interestingly, the self-assembly of different RBD-LS fusion proteins can produce multiplexed nanoparticles displaying a panel of different RBD variants in multiple copies onto the same NP scaffold. We have already demonstrated that a bivalent mRNA vaccine encoding for the wild type and the alpha (B.1.1.7) variant RBD-NP elicits the same anti-RBD neutralizing antibody titers as the monovalent mRNA RBD-NP vaccine (Fig. 2 B).

As a large share of the population in many countries is already vaccinated with mRNA-based vaccines there is increased interest for heterologous prime/boost vaccination for the control of emerging SARS-CoV-2 variants. Optimal antibody responses are induced by heterologous regimens involving priming with mRNA and boosting with nanoparticles (Vogt et al., 2021). Thus, the mRNA RBD-NP platform is suitable for an extension of current mRNA-based vaccine regimens.

A convergence of acquired RBD mutations was observed in five residues (417, 452, 478, 484 and 501) for the Pango lineages designated variants of concern by the WHO, B.1.1.7 (alpha), B.1.351 (beta), P.1 (gamma) and B.1.617.2 (delta). Global SARS-CoV-2 RBD surveillance (GISAID EpiCoV™ database) will indicate in the coming months any similar convergence of RBD mutations in a limited number of residues in the sublineages BA.1, BA.1.1, BA.2 and BA.3 of the omicron variant. Results of an ongoing study of a mRNA vaccine using the same RBD-NP platform against the omicron variant of SARS-CoV-2 will guide the design of a multivalent mRNA RBD-NP vaccine effective against foreseeable SARS-CoV-2 variants of concerns.

## ACKNOWLEDGMENTS

We are grateful to Etienne Simon-Loriere (Institut Pasteur, Paris, France) for helping in the design of in vivo challenge experiment and to Matthieu Prot (Institut Pasteur) for producing the SARS-CoV-2 delta variant stock and inoculum. We thank Dr. Gert Zimmer and Dr. Obdulio Garcia (Institute of Virology and Immunology and University of Bern, Switzerland) for assistance on SARS-CoV-2 based neutralization assays. We thank Dr. Andrew Ward (The Scripps Research Institute, La Jolla, CA, USA) for assistance on electron microscopy. We thank Dr. Anton Maximov (The Scripps Research Institute, La Jolla, CA, USA) for providing a pHL-sec plasmid and Pr. E. Yvonne Jones (Welcome Centre for Human Genetics, University of Oxford, UK) for agreeing to transfer this plasmid for cloning and transient expression of RBD-NP. We thank Dr. Sujay Singh and Pershing Billings (Abeomics, San Diego, CA, USA) for assistance on RBD-NP transient expression. Constructs for mRNA transcription were generated and cloned by Eton Bioscience in San Diego, CA, USA. VT received support for the Swiss National Science Foundation as a part of the NCCR RNA & Disease (grant 182880), a National Centre of Competence (or Excellence) in Research. AA is supported by amfAR Mathilde Krim Fellowships in Biomedical Research no. 110182-69-RKVA.

## AUTHOR CONTRIBUTIONS

PB conceived, designed, supervised and analyzed the experiments and wrote the paper; XM, LC designed, supervised, performed and analyzed the mouse challenge and protection experiments; IM designed and performed the formulation; VT, GTB supervised, performed and analyzed the SARS-CoV-2 delta neutralization tests; NS performed statistical analysis and edited the manuscript; BT cloned the constructs for SARS-CoV-2 delta variant; AD performed and analyzed *in vitro* assays; AA performed and analyzed electron microscopy; GS, HKL performed and analyzed protein expression and purification; MV provided formulation material; AS supervised RNA *in vitro* transcription and purification; JH assisted with study design and supervision and reviewed the manuscript.

## DECLARATION OF INTERESTS

PB and JH are the founders and owners of Phylex BioSciences, Inc., the company that holds IP related to RBD-NP. PB is a named inventor on several RBD-NP vaccine patents. The other authors declare no commercial or financial conflict of interest.

## MATERIALS AND METHODS

### Ethics statement

Animal welfare for the mouse studies in the U.S. was conducted in compliance with the U.S. Department of Agriculture (USDA) Animal Welfare Act (9 CFR Parts 1, 2, and 3) and the Guide for the Care and Use of Laboratory Animals, 8^th^ Edition, National Academy Press, 2011, Washington, DC, USA. The studies design and animal usage were reviewed and approved by the Institutional Animal Care and Use Committee under numbers 20-024, 21-006, 21-020 and 21-029 for compliance with regulations prior to study initiation.

Infectious challenge of immunized mice was carried out at the Institut Pasteur in compliance with French and European regulations (Directive 2010/63EU). Experimental protocols were approved by the Institut Pasteur Ethics Committee under number dap210050 and authorized by the French Ministry of Research under number 31816.

### Mice

CB6F1/J mice and B6.Cg-Tg(K18-ACE2)2Prlmn/J transgenic mice were purchased from The Jackson Laboratory (SN 100007 and 034860, respectively) under a license agreement. During the immunization period, mice were kept in individually ventilated cages in the animal facility of BTS Research in San Diego, CA, USA. Infectious challenge was performed in isolators, in a BSL-3 mouse facility at the Institut Pasteur in Paris, France. Mice were fed standard chow ad libitum and kept in a 14:10 light:dark cycle.

### Transient expression of RBD-NP protein

FreeStyle 293 cells (2.10^6^) in 500 ml of SFM media were transfected using 600 μg of pHL-sec plasmid DNA containing RBD-NP cDNA for transient expression of glycosylated RBD-NP according to the manufacturer’s protocol (Life Technology). After 7 days of transfection cells were harvested, spun down at 4600 g for 30 min at 4°C. The supernatant was filtered through a 0.22 μm filter and was stored at 4°C until ready for next step. For the purification of RBD-NP, 5 ml of lectin beads (*Galanthus nivalis* lectin (GNL)-agarose beads (Vector Labs: AL-1243-5) were added to the filtered transfected supernatant and rocked overnight at 4°C. The supernatant and bead mixtures were loaded onto a 2 ml disposable column and was allowed to flow through via gravity. The column was washed with PBS and the bound glycoproteins were eluted with glycoprotein eluting solution for mannose/glucose binding lectins (Vector Labs: ES-1100). The glycoproteins were dialyzed in PBS, concentrated, and were analyzed by SDS-PAGE, HPLC and western blot. For the western blot analysis, the samples were run on SDS-PAGE, transferred to a PVDF membrane. The membrane was incubated with a polyclonal anti-RBD antibody (Sino Biological: 40592-T62) at 1:2000 dilution followed by incubation with HRP conjugated anti-rabbit polyclonal antibody and reaction was detected by chemiluminescence and revealed the expressed protein at the estimated size of the RBD-NP.

### Electron microscopy of RBD-NP

A 15μl sample of expressed protein at 1mg/ml concentration was diluted in TBS at 20μg/ml and applied to a carbon coated copper grid (440-mesh) with uranyl-formate staining. Grids were prepared in duplicates and EM imaging was performed with a TF20 microscope (200kV) and Tietz 4k/4k camera. 186 micrographs were acquired and 3271 particles were picked up manually and 2D- aligned into 20 classes for analysis.

### mRNA encoding RBD-NP

mRNA encoding the RBDdelta fused with LS was designed with a minimal 5’ UTR of 14 nucleotides, a human beta-globin 3’UTR, 3’ polyadenylation with a poly(A) tail of 80 nucleotides, and unmodified nucleosides.

### In vitro transcription of mRNA

A construct with the 5’ minimal untranslated region, the human IL-2 signal sequence, a nucleotide sequence encoding RBDdelta fused with LS, the human beta-globin 3’UTR, a poly(A) tail of 80 adenosine residues and the BsmBI restriction site was cloned into a pUC19 vector. The supercoiled pUC19 DNA was upscaled, linearized with BsmBI, and purified. In vitro transcription was performed with T7 RNA polymerase in a 2mL reaction. The mRNA was capped with a cap 1 structure on the 5’ end by vaccinia 2’-*O*-methyltransferase enzymatic capping. Capped mRNA was purified by reverse phase chromatography using POROS™ resin (Thermo Fisher) followed by tangential flow filtration (TFF). Final yield of purified mRNA was 2.88 g/l of IVT reaction. RNA integrity was analyzed by denaturing PAGE. mRNA sample and ladder were denatured at 70°C for 2 minutes and kept on ice, then 200ng of mRNA mixed with diluent marker was loaded on gel.

### Formulation

The mRNA was complexed by addition of clinical grade protamine to the mRNA at a mass ratio of 1:5. The vaccine was prepared on each injection day with final total mRNA concentration of 840mg/l. mRNA integrity prior to and after formulation was analyzed on 1% agarose gel with SYBR safe stain (Thermo Fisher).

### Immunization

All groups of immunized mice were primed at week 0 and boosted at week 3 with the same dose of 42µg/50µl by intramuscular injection at the caudal thigh under 1- 5% isoflurane anesthesia with a needle-free injection system (Tropis® injector prototype modified for mouse injection, PharmaJet). Blood was collected by retro-orbital sinus or submandibular vein puncture and serum samples were prepared in week 0 (prior to prime), 3 (prior to boost), 6, and additionally in week 8 (prior to viral challenge) for the transgenic mice. As an exception for the injection route study (Fig. 2 A) boost was at week 2 and blood collection at week 4.

### Viral and infectious challenge

The Delta variant (B.1.617.2) strain hCoV-19/France/HDF-IPP11602i/2021 (GISAID ID: EPI_ISL_3030060) was supplied by the National Reference Centre for Respiratory Viruses hosted by Institut Pasteur (Paris, France) and headed by Pr. Sylvie van der Werf. The human sample from which strain hCoV- 19/France/HDF-IPP11602i/2021 was isolated has been provided by Dr Raphaël Guiheneuf, CH Simone Veil, Beauvais France. Moreover, the strain hCoV- 19/France/HDF-IPP11602i/2021 was supplied through the European Virus Archive goes Global (Evag) platform, a project that has received funding from the European Union’s Horizon 2020 research and innovation programme under grant agreement No 653316.

At week 8 post prime, all transgenic mice were anesthetized (ketamine/xylazine) and inoculated with an intranasal dose of 10^4^ plaque-forming units (PFU) of SARS-CoV-2 delta variant (B.1.617.2) in a 40µl volume. Eight immunized mice and 8 unvaccinated controls were followed until 14 days post-infection (dpi) for body weight variation and clinical signs of COVID-19 disease. Each of four criteria (ruffled fur, hunched posture, reduced locomotion and difficult breathing) received a 0-3 score and were added into a global score. Mice with 25% body weight loss or a score of 8 were euthanized.

### Viral load and viral titer in lungs

After infection 8 immunized mice and 8 unvaccinated controls were euthanized at 3 dpi for measurement of viral load and viral titer in lung homogenates. Viral RNA was extracted using an extraction robot IDEAL-32(IDsolutions) and the NucleoMag Pathogen extraction kit (Macherey Nagel). Viral RNA quantification was performed by quantitative reverse transcription PCR (RT-qPCR) using the IP4 set of primers and probe (nCoV_IP4-14059Fw GGTAACTGGTATGATTTCG and nCoV_IP4- 14146Rv CTGGTCAAGGTTAATATAGG giving a 107bp product, and nCoV_IP4-14084Probe(+) TCATACAAACCACGCCAGG [5’]Hex [3’]BHQ-1 19) and the Luna Universal Probe One-Step RT-qPCR Kit (NEB). Serial dilutions of a titrated viral stock were analyzed simultaneously to express viral loads as eqPFU per gram of tissue.

For plaque assay, 10-fold serial dilutions of samples in DMEM were added onto VeroE6 monolayers in 24 well plates. After one-hour incubation at 37°C, the inoculum was replaced with equivalent volume of 5% FBS DMEM and 2% carboxymethylcellulose. Three days later, cells were fixed with 4% formaldehyde, followed by staining with 1% crystal violet to visualize the plaques.

### Inflammatory cytokine and chemokine content

Total RNA was prepared from one lung lobe collected at 3 dpi using lysing matrix D (MP Biomedical) containing 1 ml of TRIzol reagent (Ambion, Life technologies) and homogenization at 20 s at 4.0 m/s twice using MP Biomedical Fastprep 24 Tissue Homogenizer. cDNA was synthesized from 200ng of RNA in the presence of 2.5 µM of Random Hexamer primers, 5mM of each deoxyribonucleotides, 40U of RNase Inhibitor and 200U of SuperScript II Reverse Transcriptase (Thermo Fisher Scientific) in 20 µL reaction. The real-time PCR was performed on QuantStudio 7 Flex Real-Time PCR System (Thermo Fisher Scientific). Reactions were performed in duplicates in a final reaction volume of 20 µL containing 10 µL of Power SYBR Green Master Mix (Applied Biosystems), 2 µL of cDNA and 0.6 µL of each forward and reverse primers at a final concentration of 300nM (Table XX). The following thermal profile was used: a single cycle of polymerase activation for 2 min at 50°C and 10 s at 95°c, followed by 40 amplification cycles of 15 s at 95°C and 1 min 60°C (annealing-extension step). Mouse *Gapdh* was used as an endogenous reference control to normalize differences in the amount of input nucleic acid. The average CT values were calculated from the technical replicates for relative quantification of target cytokines/chemokines. The differences in the CT cytokines/chemokines amplicons and the CT of the endogenous reference control, termed dCT, were calculated to normalize for differences in the quantity of nucleic acid. Relative expression was calculated as 2^dCT^.

### SARS-CoV-2 delta neutralizing antibody detection assay

Sera samples of week 0, 3, 6 and 8 were analyzed with the cPass™ SARS-CoV- 2 neutralization antibody detection kit (GenScript, Cat #L00847) to detect any antibodies that neutralize the interaction between the RBDdelta and the ACE2 receptor. The kit contains two key components: the horseradish peroxidase (HRP) conjugated recombinant SARS-CoV-2 RBD (HRP-RBD), and a capture plate coated with human ACE2 receptor protein (hACE2). The protein-protein interaction between HRP-RBD and hACE2 is blocked by neutralizing antibodies against the RBD. The assay was performed as per the manufacturer’s protocol, except the HRP-RBD component was replaced with HRP conjugated RBDdelta. First, the samples and controls were pre-incubated with the HRP-RBDdelta for 30 min at 37°C to allow the binding of the circulating neutralization antibodies to HRP-RBDdelta. The mixture was then added to the capture plate and incubated for 15 min at 37°C. The unbound HRP-RBDdelta as well as any HRP-RBDdelta bound to non-neutralizing antibody was captured on the plate, while the circulating neutralization antibodies- HRP-RBDdelta complexes remained in the supernatant and were removed during washing. After washing steps, 3, 3’, 5, 5’-tetramethylbenzidine (TMB) solution was added and the plate was incubated in dark at 20°C for 15 min with the color turning blue. The reaction was quenched by adding a stop solution, with the color turning yellow, and the final solution was read immediately at 450 nm in a microtiter plate reader. The absorbance of the sample was inversely dependent on the titer of the anti-SARS-CoV-2 neutralizing antibodies. The RBDdelta-ACE2 interaction inhibition rate was calculated with the net optical density (OD) of sample and negative control with the formula: Inhibition = (1 – OD value of sample/OD value of negative control) x 100%. Neutralizing antibodies were detected with a cutoff inhibition value of 30% established by the manufacturer in a human clinical study with a combined cohort of healthy individuals (n=88) and SARS-CoV-2 positive patients confirmed by RT-PCR (n=26).

### Generation of infectious cDNA clone using TAR cloning and rescue of recombinant delta spike virus (SARS-CoV-2^S-delta^)

To introduce the variant specific mutations into the spike gene, we used twelve 50 bp primers containing the desired nucleotide changes in a PCR reaction. These PCR products were used to replace WU-Fragments 9 and 10 (covering the spike region). The WU-Fragments encoding the whole SARS-CoV-2 genome are described in Thao et al. (2020). The WU-Fg. 1.3-8, 11, 12 and the newly created PCR fragments with 50 bp homologous overlaps, were then used for the in-yeast TAR cloning method as described previously (Thao et al., 2020) to generate infectious cDNA clones. In vitro transcription was performed for EagI-cleaved YACs and PCR-amplified SARS-CoV-2 N gene using the T7 RiboMAX Large Scale RNA production system (Promega) as described previously. Transcribed capped mRNA was electroporated into baby hamster kidney (BHK-21) cells expressing SARS-CoV-2 N protein. Electroporated cells were co-cultured with susceptible VeroE6 cells expressing TMPRSS2 to produce passage 0 (P.0) of the recombinant SARS-CoV-2^S-delta^. Subsequently, progeny virus was used to infect fresh VeroE6-TMPRSS2 cells to generate P.1 stocks for downstream experiments.

### SARS-CoV-2 delta neutralization assay

Vero-E6 cells were seeded at a density of 1.5x10^4^ cells/100μl per well (in DMEM supplemented with 10% FBS, 1% non-essential amino acids (NEAA) and 1% Penicillin-Streptomycin) in 96-well cell culture plates one day before and incubated over night at 37°C, 5% CO2. Two-fold dilution series of control sera and serum samples of week 3, 6 and 8 were prepared in quadruplicates in 96-well cell culture plates using unsupplemented DMEM cell culture medium (50μl /well). To each well, 50μl of DMEM containing 100 plaque forming units (PFU) of SARS-CoV-2^S-delta^ from P.1 stock described above were added and incubated for 60 min at 37°C. Subsequently 100 μl of serum and virus mixtures were added on confluent Vero E6 cells and 96-well plates were incubated for 72 h at 37°C. The cells were fixed for 10 min at room temperature with 4% buffered formalin solution, then cells were counterstained with a solution containing 1% crystal violet for another 10 min at room temperature. Finally, the microtiter plates were rinsed with deionized water and immune serum-mediated protection from cytopathic effect was visually assessed.

### Statistical analysis

Statistical analyses were performed using Prism v9.3.1 (GraphPad Software) (Fig. 2-3, 5, 10-11) or using R Statistical Software v4.1.2 (R Core Team 2021) (Fig. 6- 9). Some datasets were analyzed after Log-transformation as indicated in the figure legends. Statistical differences between groups in datasets with one categorical variable were evaluated by two sample t-test (2 groups) or one-way ANOVA (more than 2 groups) corrected for multiple comparisons unless indicated otherwise. Correlations between two variables were evaluated with simple linear regression and Pearson correlation coefficient unless specified otherwise. *p* values ≤ 0.05 were considered significant with the following reporting style in the figures:

ns *p* > 0.05

* p ≤ 0.05

** p ≤ 0.01

*** p ≤ 0.001

**** p ≤ 0.0001

## Notes

### Competing Interest Statement

Pascal Brandys and Jens Herold are the founders and owners of Phylex BioSciences, Inc., the company that holds IP related to RBD-NP. Pascal Brandys is a named inventor on several RBD-NP vaccine patents. The other authors declare no commercial or financial conflict of interest.

